# Discontinuous transcription of ribosomal DNA in human cells

**DOI:** 10.1101/768929

**Authors:** Evgeny Smirnov, Peter Trošan, João Victor de Sousa Cabral, Pavel Studený, Sami Kereïche, Kateřina Jirsová, Dušan Cmarko

## Abstract

Numerous studies show that various genes in all kinds of organisms are transcribed discontinuously, i.e. in short bursts or pulses with periods of inactivity between them. But it remains unclear whether ribosomal DNA (rDNA), represented by multiple copies in every cell, is also expressed in such manner. In this work, we synchronized the pol I activity in the populations of tumour derived as well as normal human cells by cold block and release. Then using special software for analysis of the microscopic images, we measured the intensity of transcription signal revealed by incorporated 5-fluorouridine (FU) in the nucleoli at different time points after the release. We found that the ribosomal genes in the human cells are transcribed discontinuously with periods ranging from 45 min to 75 min. Our data indicate that the dynamics of rDNA transcription follows the undulating pattern, in which the bursts are alternated by periods of rare transcription events.

## Introduction

Numerous studies show that genes in all kinds of organisms, from prokaryotes to mammals, can be transcribed in short bursts or pulses alternated by periods of silence (reviewed in Smirnov et al. [1]) The probability of such mode of expression was suggested long ago;[2] now it seems that the discontinuous transcription is a common feature of the gene expression, at least in mammalian cells.[3–12] The periodical switches of the promoter between the active and “refractory” states may be crucial in the efficient regulation of the gene expression.[13–17] General considerations suggest even more significant role of the phenomenon in the dynamic organization of the cell, since the pulsing mode of one process is likely to be a cause and a consequence of pulsing in other processes. Thus, RNA processing, which is closely linked to the RNA synthesis, seems to be discontinuous.[9] A spontaneous heterogeneity of gene expression occasioned by transcriptional fluctuations may influence cell behaviour in changing environmental conditions and in the course of differentiation.[18]

The discontinuous character of transcription has been detected by various methods (reviewed in Smirnov et al. [1]) The number of transcripts produced in a certain (sufficiently short) period of time may be determined with high precision by single molecule RNA fluorescence in situ hybridisation (smFISH).[19–21] The results of such quantification alone provide indirect, but valuable information for modelling the expression kinetics in a cell population or tissue, when the studied gene is supposed to be transcriptionally active in all the cells. Methods based on the allele-sensitive single-cell RNA sequencing also allow to reveal and characterize the transcription bursting.[22] To monitor gene expression in real time, cells are transfected with constructs providing a fluorescent signal that corresponds to the expression of a particular gene. In a gene trap strategy, a luciferase gene is inserted under the control of endogenous regulatory sequences. Since both the luciferase protein and its mRNA are short-lived, the method allows to calculate the key parameters of the transcriptional kinetics. Probably the most popular *in vivo* method is based on the use of bacteriophages derived fluorescent coat proteins, such as MS2 or PP7, fused with GFP, which allows to visualize a bunch of the nascent RNA molecules accumulated around one gene.[4, 23, 24]

So far, the pulse-like transcription is well documented only in the genes transcribed by RNA polymerase II. It is not clear yet whether ribosomal DNA (rDNA) is also expressed discontinuously. In human cells, the clusters of multiple rDNA repeats, known as Nucleolus Organizer Regions (NORs), are situated on the short arms of the acrocentric chromosomes. Each repeat includes a gene coding for 18S, 5.8S and 28S RNAs of the ribosomal particles and an intergenic spacer.[25–30] In the interphase nucleus the rDNA provides the basis for the formation of nucleoli. The transcription by pol I and the first steps of rRNA processing take place in the special nucleolar units (FC/DFC) composed of fibrillar centers (FC) and dense fibrillar components (DFC).[31–42] The units correspond in light microscopy to the “beads” forming nucleolar necklaces,[43–46] and each unit is believed to accommodate a single transcriptionally active gene.[33, 39, 47, 48] The intensity of the rDNA transcription is usually very high throughout the entire interphase, especially at the S and G2 phases.[49] Now most of the methods used for the detection of the transcription fluctuation are hardly applicable to the ribosomal genes, since one cell usually contains hundreds of such genes. An alternative method was designed for direct measurements of rDNA transcription in the live cells by using the label-free confocal Raman microspectrometry.[50] This work revealed an undulatory character of the ribosomal RNA production in the whole nucleoli. In our earlier study on tumour-derived cells expressing a GFP-RPA43 (a subunit of pol I) fusion protein, we have observed specific fluctuations of the fluorescence signal in the individual FC/DFC units.[51] We also found high correlation of pol I and incorporated FU signals within the units. These data suggested that the ribosomal genes are transcribed in a pulse-like manner.

In the present work we used a different approach to the study of the discontinuous transcription of ribosomal genes in human cells. Namely, we synchronized the pol I activity in the cell population by cold block and release. Then, using specially designed software we measured the intensity of transcription signal, incorporated 5-fluorouridine (FU), in the nucleoli and individual FC/DFC units at different periods after the release. This enabled us to detect transcription fluctuations of ribosomal genes in tumour derived as well as normal human cells and to reveal special properties of this fluctuation.

## Methods

### Ethics

The study followed the standards of the Ethics Committees of the General Teaching Hospital and the First Faculty of Medicine of Charles University, Prague, Czech Republic (Ethics Committee of the General University Hospital, Prague approval no. 8/14 held on January 23, 2014), and adhered to the tenets set out in the Declaration of Helsinki. We obtained human cadaver corneoscleral rims from 10 donors, which were surplus from surgery and stored in Eusol-C (Alchimia, Padova, Italy), from the Department of Ophthalmology, General University Hospital in Prague, Czech Republic, for the study. On the use of the corneoscleral rims, based on Czech legislation on specific health services (Law Act No. 372/2011 Coll.), informed consent is not required if the presented data are anonymized in the form.

### Cell cultures

Human limbal epithelial cells (LECs) were obtained from XY cadaver corneoscleral rims after cornea grafting at University Hospital Kralovske Vinohrady, Prague, Czech Republic. The mean donor age ± standard deviation (SD) was 63.5 ± 6.5 years. Tissue was stored in Eusol-C (Alchimia, srl., Ponte San Nicolò, Italy) preservation medium at +4°C. The mean storage time ± SD (from tissue collection until explantation) was 7.2 ± 3.6 days. The corneoscleral rims were prepared as described before.[52, 53] Shortly, corneoscleral rims were cut into 12 pieces and placed in a 24-well plate (TPP Techno Plastic Products AG, Trasadingen, Switzerland) on Thermanox plastic coverslips (Nunc, Thermo Fisher Scientific, Rochester, NY, USA). Explants were cultured in 1 ml of complete medium [1:1 DMEM/F12, 10% FBS, 1% AA, 10 ng/ml recombinant EGF, 0.5% insulin-transferrin-selenium (Thermo Fisher Scientific), 5 μg/ml hydrocortisone, 10 μg/ml adenine hydrochloride and 10 ng/ml cholera toxin (Sigma-Aldrich, Darmstadt, Germany)]. The culture media were changed every 2 – 3 days until the cells were 90–100% confluent (after 2-4 weeks).

HeLa cells were cultivated at 37°C in Dulbecco modified Eagle’s medium (DMEM, Sigma) containing 10% fetal calf serum, 1% glutamine, 0.1% gentamicin, and 0.85g/l NaHCO_3_ in standard incubators.

### Plasmids and transfection

The GFP-RPA43 and GFP-fibrillarin vectors were received from Laboratory of Receptor Biology and Gene Expression Bethesda, MD.[54] The constructs were transfected into HeLa cells using Fugene (Qiagen).

### Labeling of the transcription sites

For visualization of the transcription sites, sub-confluent cells were incubated for 5 min with 5-fluorouridine (FU) (Sigma). The cells were fixed in pure methanol at −20°C and processed for FU immunocytochemistry. Incorporated FU signal was visualized using a mouse monoclonal anti-BrdU antibody (Sigma).

### Light microscopy

Confocal images were acquired by means of SP5 (Leica) confocal laser scanning microscope equipped with a 63×/1.4NA oil immersion objective. For *in vivo* cell imaging we used a spinning disk confocal system based on Olympus IX81 microscope equipped with Olympus UPlanSApo 100×/1.4NA oil immersion objective, CSU-X spinning disk module (Yokogawa) and Ixon Ultra EMCCD camera (Andor). The live cells were maintained in glass bottom Petri dishes (MatTek) at 37°C and 5% CO_2_ within a microscope incubator (Okolab).

### Software and data analysis

For measurement and counting of the transcription and other signals corresponding to individual FC/DFC units in 3D confocal images, we developed a MatLab based software.[51] The program identifies each unit by creating a maximum intensity projection of the confocal stack and blurring the projection with a Gaussian filter (σ = 8–10 pixels), defining the blurred image with a value obtained by Otsu’s method for automatic threshold selection. After that, the optical section whereupon the unit had maximum intensity was identified. The final result contains 3D coordinates of each unit, its size (full-width half-maximum), the value of χ^2^, and integral intensities in the spheres with radii 1.0, 1.5, 2.0, 2.5, 3.0, 3.5, and 4.0 pixels. The values corresponding to 1.5 pixels seemed to be the most resistant to noise and were used for presentation of the data. FC/DFC units were counted after deconvolution with Huygens software.

For measuring signals in the entire nucleoli we used a custom ImageJ plugin available at https://github.com/vmodrosedem/segmentation-correlation.[45] Based on the confocal stacks, the program identifies the regions occupied by the cell nuclei as well as nucleoli, measures their areas (in pixels), and the intensities, both integral and average, of the signal within these areas.

## Results

### 1. Effects of low temperature on the nucleolar transcription

In the control the incorporated FU is accumulated predominantly in the FC/DFC units of the nucleoli (Fig 1). The transcription signal in the nucleoplasm is of much lower intensity. After 15 min of incubation at +4**°**C (without additional supply of CO_2_, both HeLa and LECs lost the ability to incorporate 5-fluorouridin (FU)). When the cells were returned to the normal conditions (37**°**C, 5% CO_2_), transcription was partly restored in 15 min, and in 30 min the FU incorporation appeared as in the control (Fig 1).

**Fig 1.**
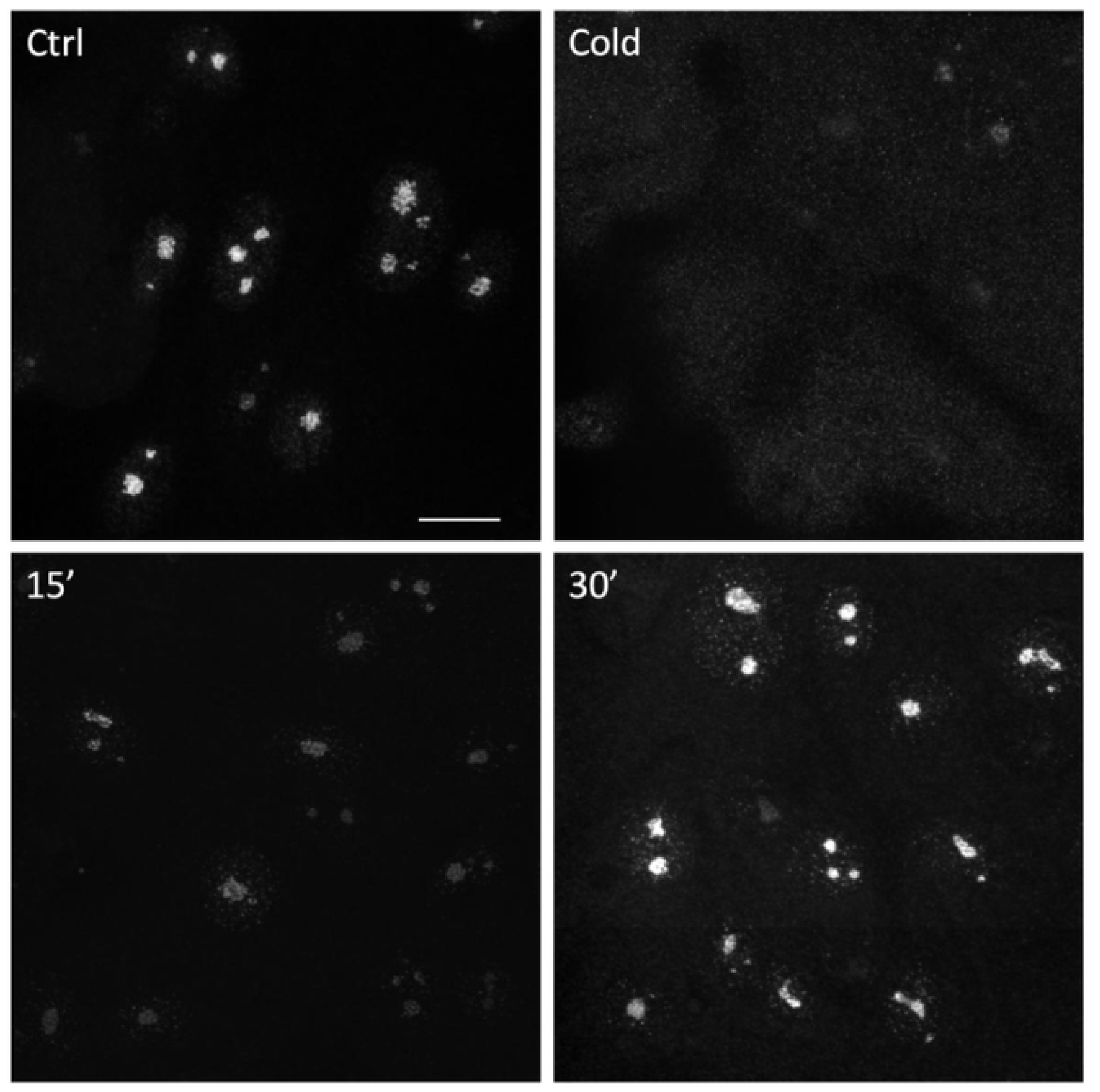
Transcription in HeLa cells is quickly inhibited at +4°C and restored at the normal conditions. The transcription signal (FU incorporation) is accumulated in the nucleoli. The signal disappeared after 15 min of cold treatment (top right); when the cells were transferred to the normal conditions, the signal was partly restored in 15 min and appeared like in the control in 30 min (bottom). Scale bar: 10µm

To assess the effect of cold on the FC/DFC units, which are the centers of rDNA transcription and early rRNA processing, we transfected the cells with GFP-RPA43 or GFP-Fibrillarin. At the low temperature the GFP-Fibrillarin signal did not change significantly, but the intensity of the RPA43 signal was decreased as average to about 60% of the control level (Fig 2). Observation of the individual cells also showed that after transferring the cells from the cold to the normal conditions, the intensity of GFP-RPA43 signal in all FC/DFC units increased, although the number of the detectable units did not change (Fig 3).

**Fig 2.**
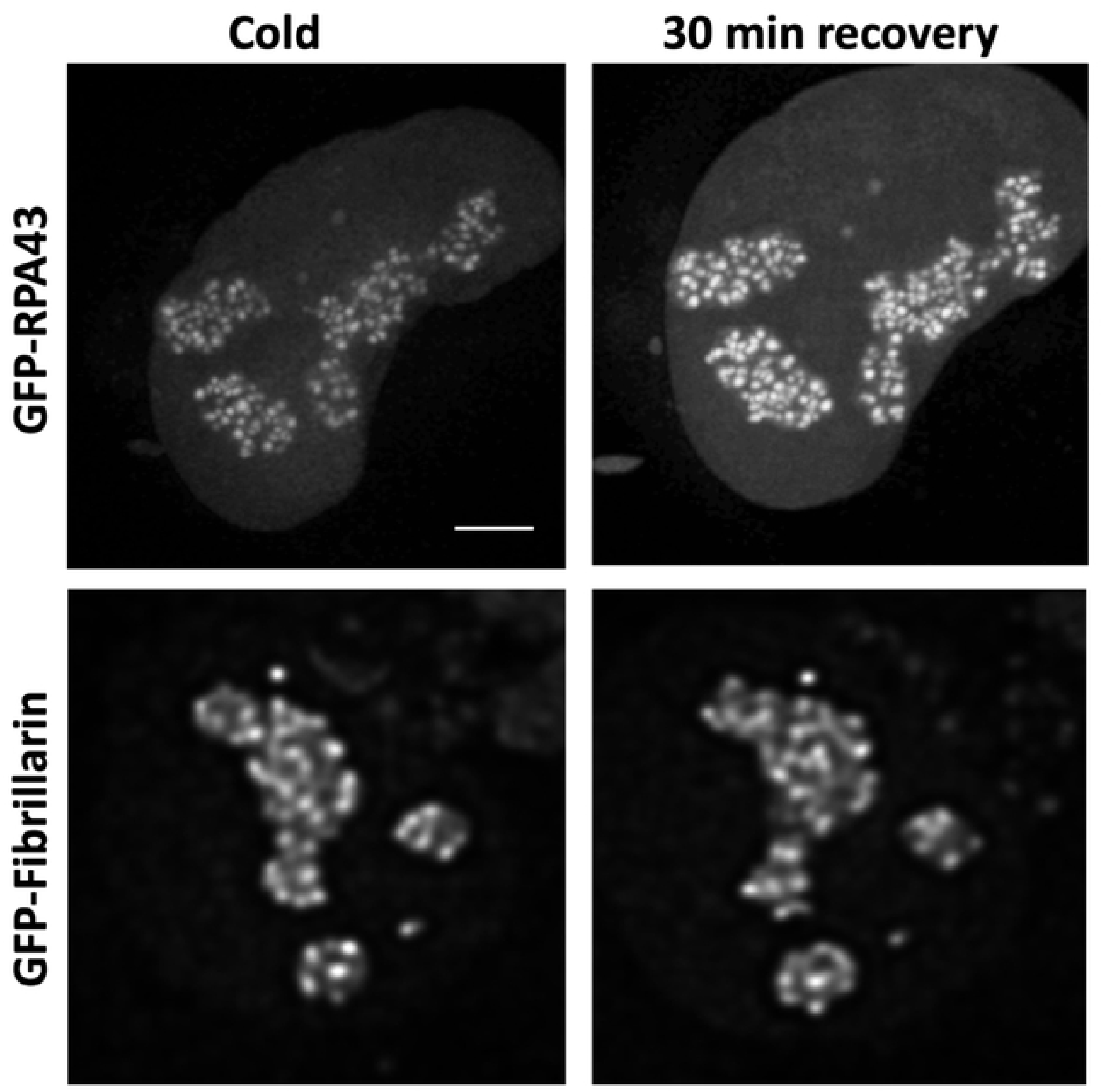
Following GFP-RPA43 and Fibrillarin-GFP signals in the transfected HeLa cells *in vivo*. The intensity of the GFP-RPA43 signal is reduced after 15min incubation at +4°C (left, top) and restored after subsequent 30 min incubation at normal conditions (top, right). The Fibrillarin-GFP signal was not significantly affected by the cooling/warming procedure (bottom). Scale bar: 5µm.

**Fig 3.**
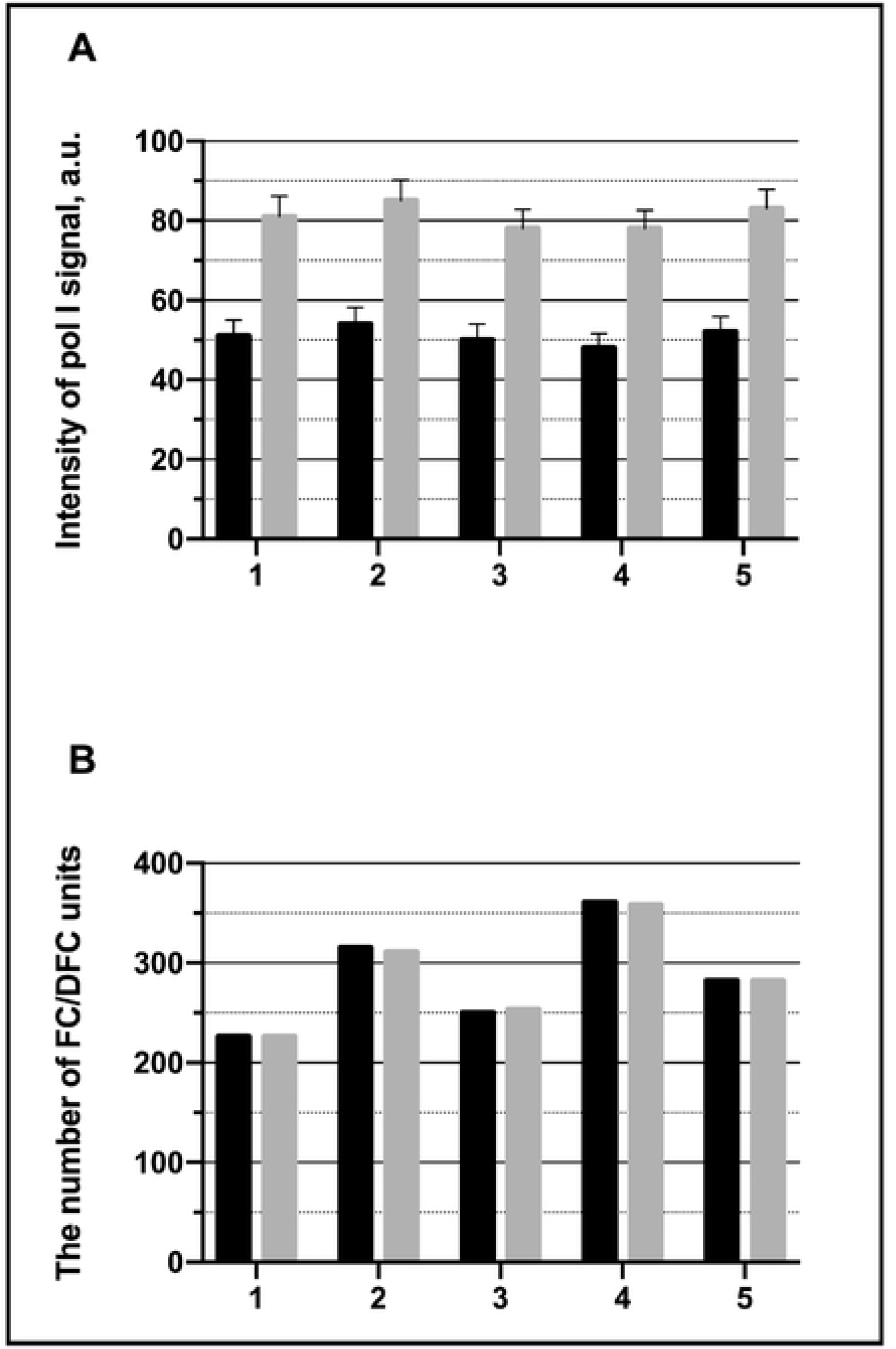
Effects of cooling/warming (as in Fig. 2, top) on the FC/DFC units *in vivo* in the transfected HeLa cells. A: intensity of the GFP-RPA43 signal in the individual units after 15min incubation at +4°C (black bars) and after subsequent 30 min incubation at 37°C (grey bars). Five cells were observed, and five selected units were followed in each cell. B: the total number of the GFP-RPA43 positive units in five cells after 15min incubation at +4°C (black bars) and after subsequent 30 min incubation at 37°C (grey bars). The experiment indicates that at the low temperature pol I escapes from the FC/DFC units.

These experiments show that low temperature causes not only quick inhibition of the rDNA transcription, but also significant though not complete depletion of the pol I pools in the nucleoli.

### 2. Synchronization of the nucleolar transcription in HeLa cells by cold treatment

The experiments described in the previous section indicate that at the low temperature the ribosomal genes are brought to a silent state with a diminished RPA-GFP signal within the FC/DFC units which implies a decreased number of pol I complexes bound to the genes. This synchronization procedure was used for the study of the discontinuous expression of the rDNA in HeLa and LEC cells. Namely, the cells were incubated in cold medium (+4**°**C) for 1 h, then transferred to the normal conditions and fixed at different time points from 15 to 150 min with the interval of 15 min. FU was added to the cultivation medium 5 min prior to each fixation. The transcription signal visualized by antibody was then measured in the nucleoli by means of the ImageJ plugin software (see Methods). The results are presented in Fig 4.

**Fig 4.**
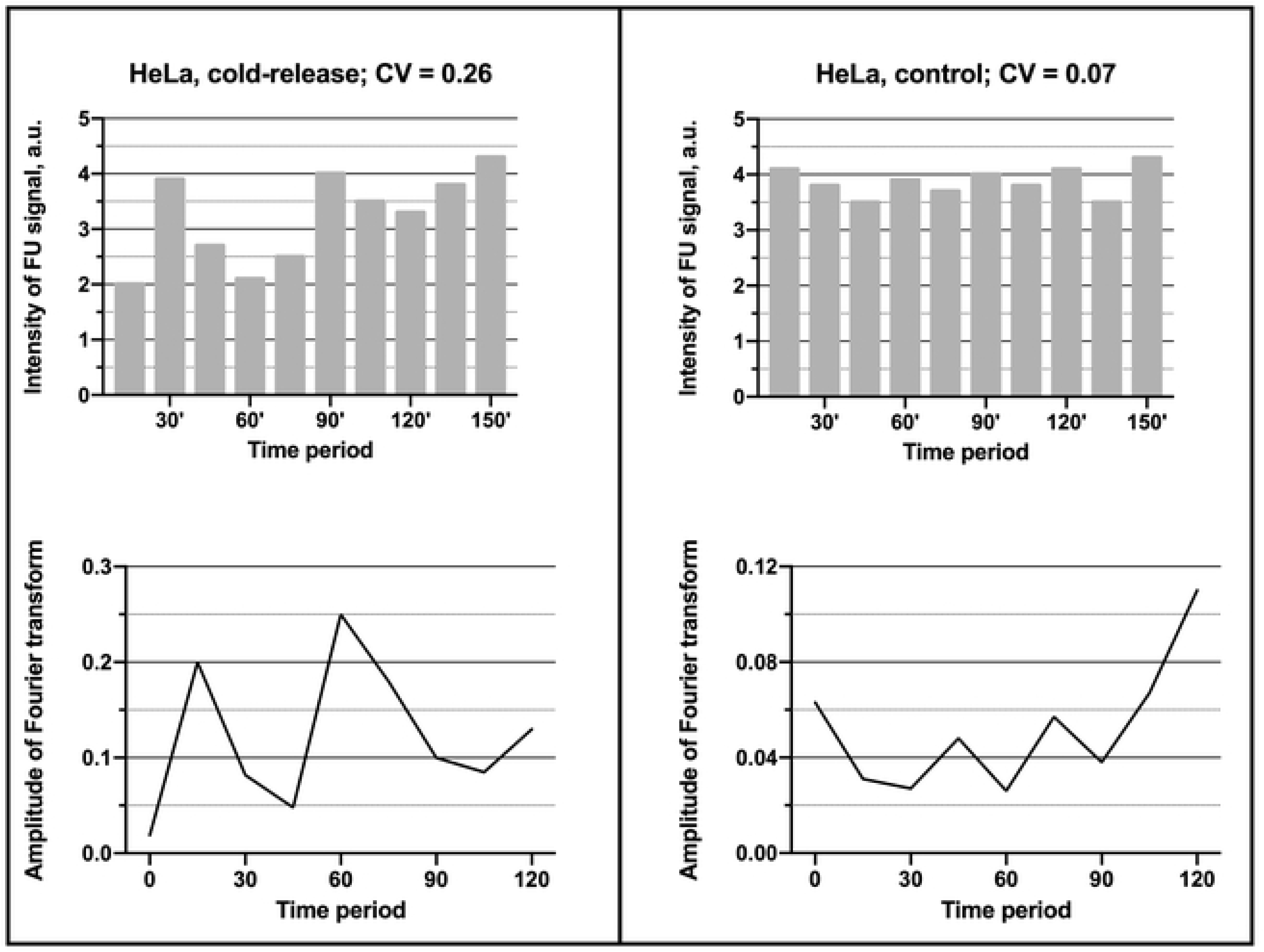
Fluctuation of the intensity of the transcription signal (incorporated FU) in the whole nucleoli of HeLa cells after release from the cold block (left) and in the control cells, i.e. without cold treatment (right). In the experiment the signal reaches maximal values at 30 min, 90 min, and 150 min. The graph shows mean values obtained from 50 cells in one experiment. Such experiment was repeated 8 times. CV-coefficient of variation. The bottom graphs show the respective periodograms for the experiment (left) and control (right) calculated as amplitudes of the Fourier transforms. The x-axis represents the period (min).

In all such experiments the intensity of the transcription signal increased during the first 30 min, then began to decrease. Altogether two cycles of rise and fall have been observed within the period of 150 min, the coefficient of variation (CV) was 0.26. The spectral analysis revealed a significant peak corresponding to the period of 60 min. Since the interval between the measurements was 15 min, the values of the period may be varying from 45 min to 75 min. An additional lower peak at 15 min probably reflected a high frequency noise. In the control, when the cells were kept at 37°C and fixed at different time points as in the experiment, the fluctuations of the transcription signal intensity were irregular. CV was only 0.07, and the periodogram had two peaks of low amplitude (compare the left and right parts of the Fig 4). In two experiments the period of observation was extended to 210 min, but between 150 and 210 min the fluctuations of the transcription signal appeared irregular like in the control (data not shown), which indicated that the synchrony in the cell population was lost.

These results showed that in HeLa cells the activity of pol I transcription machinery was synchronized by the cold treatment for the period of 150 min, but not longer.

### 3. Synchronization of the nucleolar transcription in human limbal epithelial cells by cold treatment

The same experimental procedure was applied to the LECs (Fig 5). In this case the first two cycles were more pronounced and the difference between control and experiment was more significant (compare Fig 5 and Fig 4). Otherwise, the dynamics of the transcription activity after the cold treatment proved to be similar in the studied cell lines. In the LECs, the periodogram had a more distinct peak at 60 min, but the synchronization also did not last longer than 150 min. CV was 0.29, i.e. slightly higher than in HeLa cells. It seems worth mentioning that our attempt to synchronize the transcription in human fibroblasts failed, for only a few of these cells recovered quickly enough after by the cold treatment.

**Fig 5.**
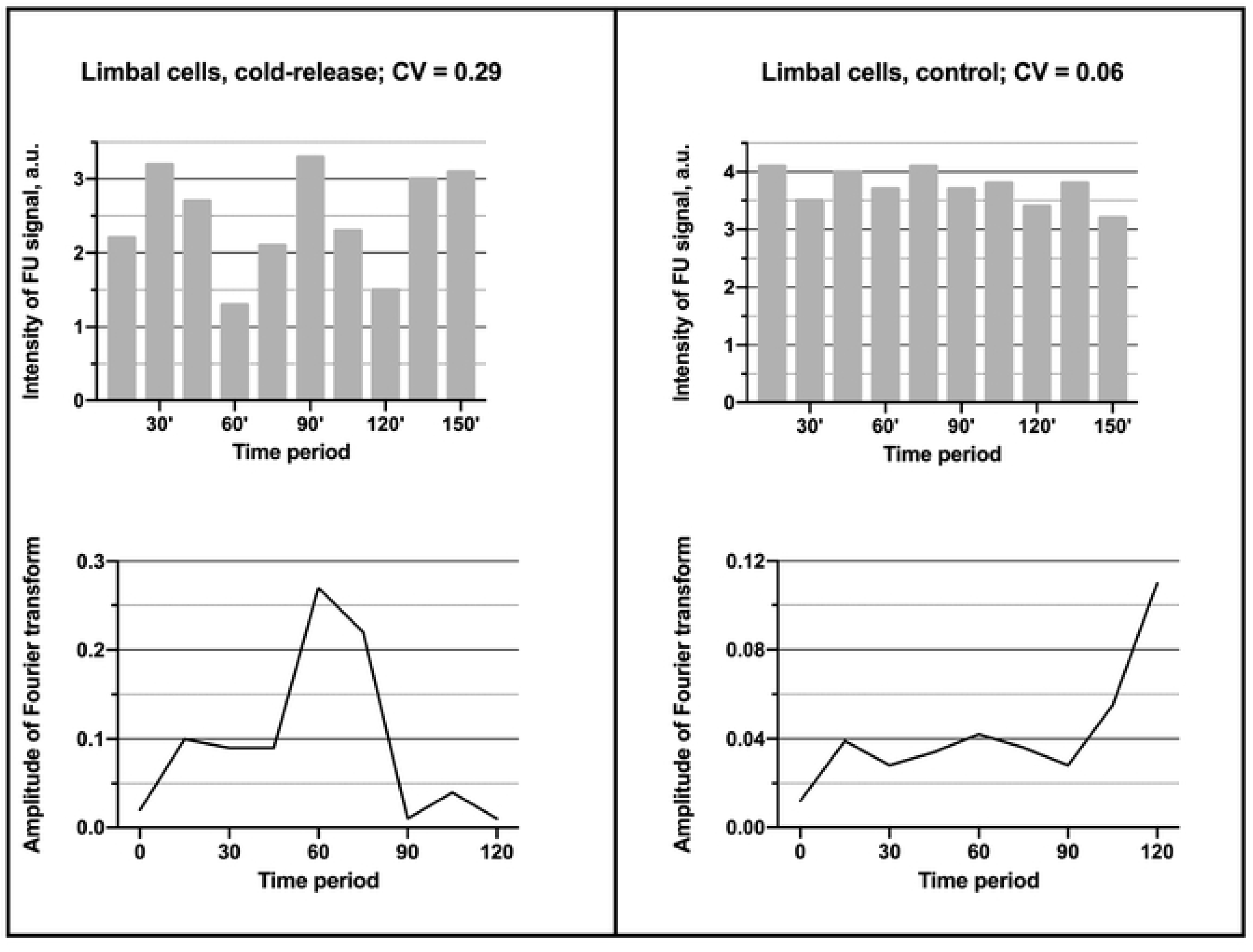
Fluctuation of the intensity of the transcription signal (incorporated FU) in the whole nucleoli of the limbal cells after release from the cold block (left) and in the control cells (right). The figure is analogous to the Fig. 4. But in this case, the undulating pattern in the experiment (top, right) is more pronounced, and the periodogram related to the experiment (bottom, left) has a more distinct peak at 60 min. The data are obtained from 8 independent experiments, and in each of them 50 cells were measured.

Thus, our experiments indicated that transcription of the ribosomal proceeds in a wave-like manner, although the employed synchronization procedure is not equally efficient in various cells.

### 4. Synchronization of the transcription in the nucleoplasm by cold treatment

When the LECs or HeLa cells were incubated at +4**°**C, the transcription ceased completely in their nucleoplasm as well as in the nucleoli. Measurement of the total FU signal after transferring the cells from the cold to the normal conditions showed symptoms of synchronization: the signal in the nucleoplasm increased for 30 min and then began to decrease (Fig. 6). The average intensity of the transcription signal in the nucleoli and nucleoplasm positively correlated, with the correlation coefficients 0.65 for the HeLa cells and 0.74 for the LECs. But, as one could expect, the total expression of the nucleoplasmic genes was less synchronized. After the initial recovery and subsequent decrease, the signal became rather noisy. The CV was 0.17 and 0.19 in the HeLa and LECs respectively. The periodograms showed a not very distinct peak at 75 min as well as a sharper peak corresponding to higher frequencies. The second peak probably reflects a noisier character of the fluctuations in the nucleoplasm as compared to the nucleoli.

**Fig 6.**
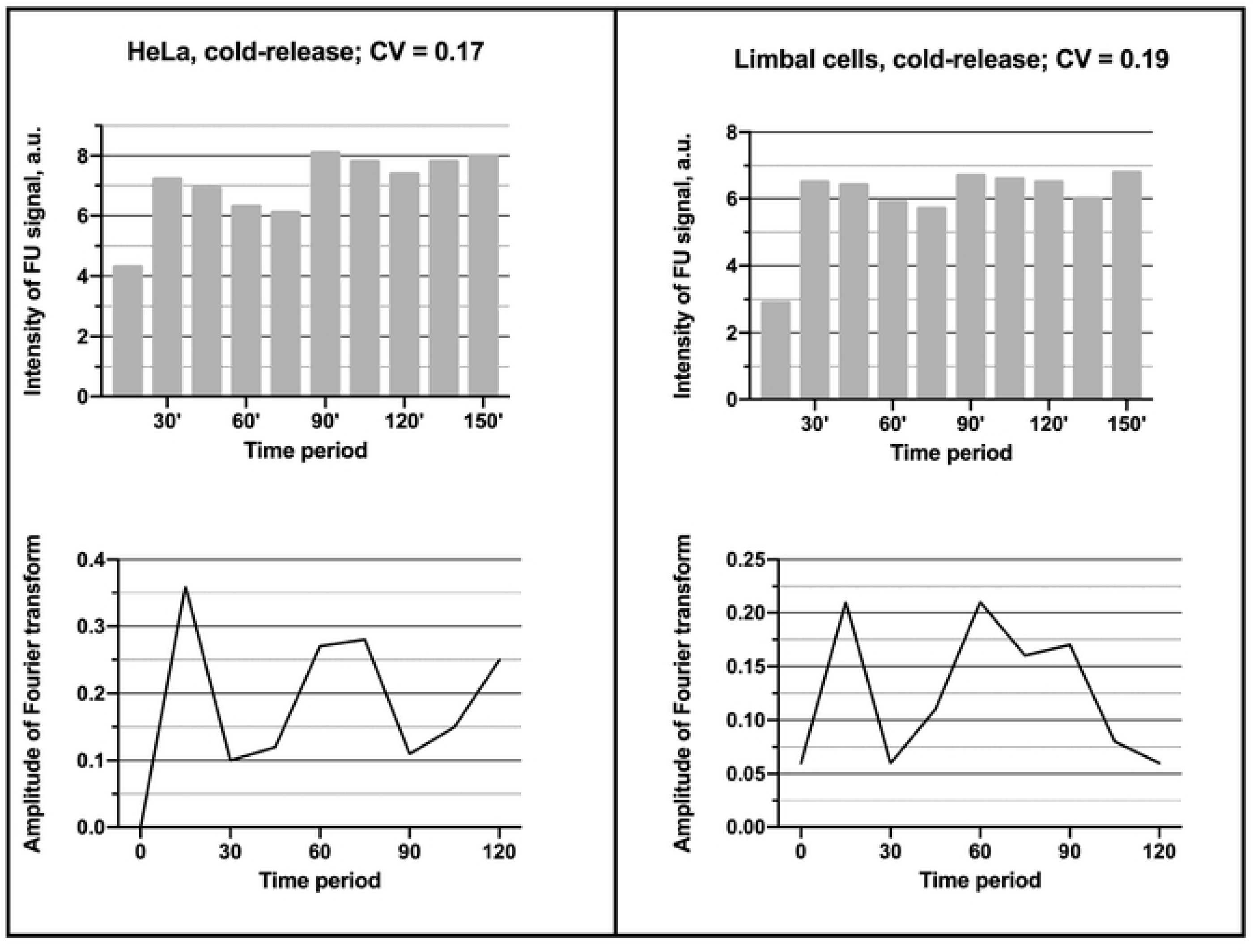
Fluctuation of the intensity of the transcription signal (incorporated FU) in the nucleoplasm of the HeLa (left, top) and limbal (right, top) cells after release from the cold block. The same cells as in Fig. 5 were used for the measurements. The periodograms related to both HeLa (left, bottom) and LECs (right, bottom) have two significant peaks.

### 5. The FC/DFC units in the course of the transcription fluctuation

According to the data presented in the sections 2 and 3 (Figs 4 and 5), the intensity of FU signal in the nucleoli at 15 min and 30 min after the cold treatment may be taken as representatives of the two extreme states of the transcriptional fluctuation in the synchronized cells. Measurement of the FU signal in the individual FC/DFC units of the LECs and HeLa cells using the MatLab based software (see Methods) showed an approximately threefold increase of the signal intensity that between 15 min and 30 min (Fig 7). But the transcription signal never disappeared from the cells completely, so that the average number of the FU-positive FC/DFC units did not change significantly (Fig 7B, right histogram).

**Fig 7.**
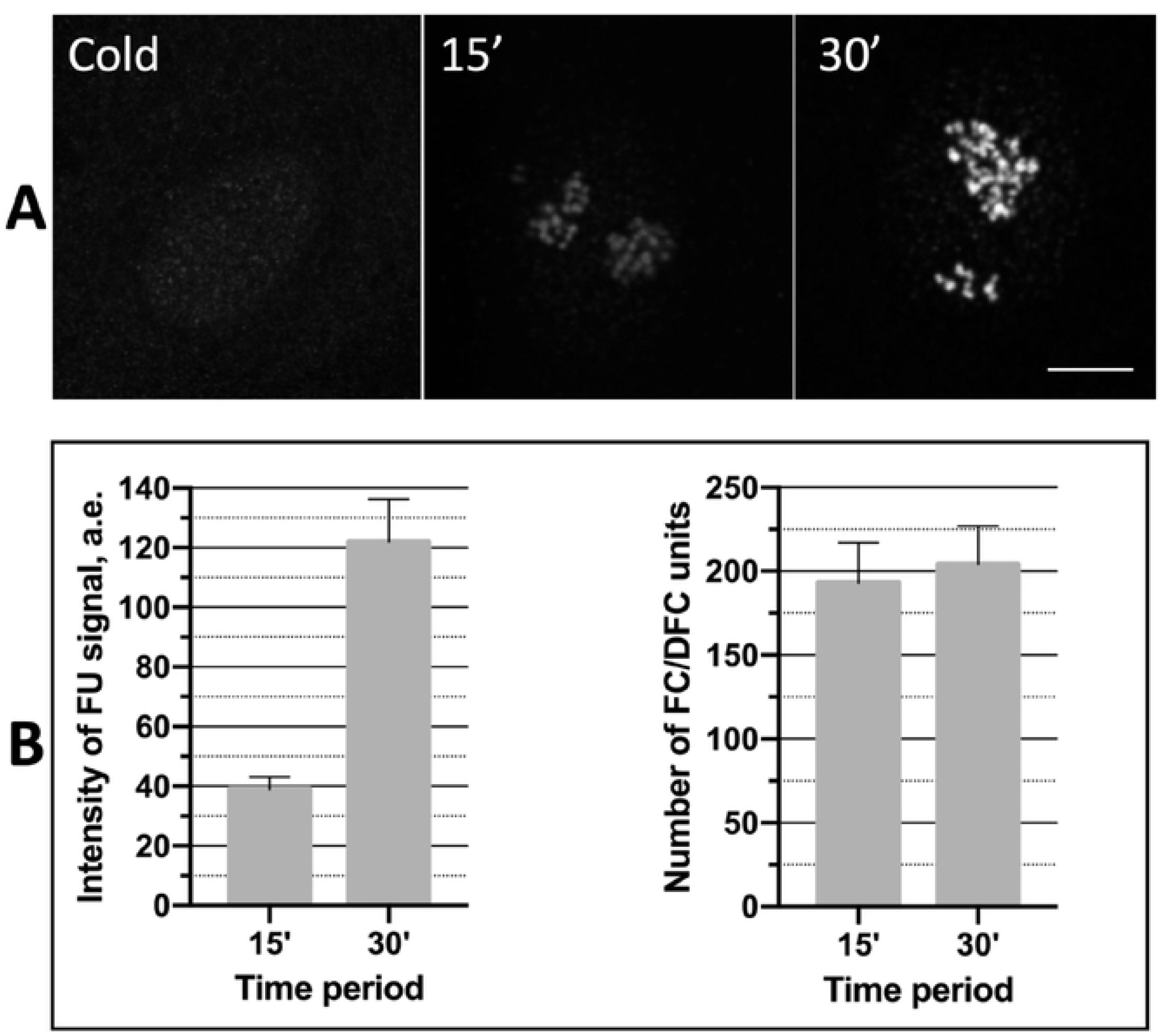
The transcription signal (incorporated FU) in the FC/DFC units of the limbal epithelial cells after the cold treatment. **A**: 15 min at +4°C, absence of the FU incorporation (left); 15 min recovery at +37°C after the cold treatment (middle); 30 min recovery at +37°C after the cold treatment (right). Scale bar: 5 µm. **B**: the average (from 50 cells) intensity of the FU signal measured in the individual FC/DFC units, 15 min and 30 min at +37°C after the cold treatment (left); the average (from 50 cells) number of the FU positive FC/DFC units, 15 min and 30 min at +37°C after the cold treatment (right). The time points 15 min and 30 min correspond respectively to state of maximal and minimal transcription intensity in the synchronized cells (see Fig 5).

## Discussion

In our experiments, when the human derived cells were incubated at +4°C, transcription in their nuclei seemed to be arrested completely (Figs 1 and 7). At the same time the pol I signal in the FC/DFC units of the nucleoli was significantly reduced (Figs 2 and 3), whereas the amount of fibrillarin, which is an essential component of the early rRNA processing, did not change significantly (Fig 2). On the other hand, previous studies, including our own, indicate that the mobile fraction of pol I, apparently responsible for the actual transcription, constitutes less than a half of the entire pool of the enzyme in the units.[51, 54] Therefore, in all probability, the pol I complexes do not “freeze” on their matrices after the arrest of the transcription by the chill shock, but rather detach themselves and escape from the units. After returning to normal conditions, the pools of the enzyme are swiftly restored, and the rRNA synthesis in the cells is synchronized. This effect was used in our work for detection of the pulse-like transcription.

In thus synchronized HeLa and LEC cells, we observed a wave-like modification of the nucleolar transcription signal with two successive peaks (Figs 4 and 5). In both kinds of cells, the predominant fluctuation period estimated by the spectral analysis was about 60 min. A similar value of the period was obtained in our previous work for the fluctuations of the GFP-RPA43 signal.[51] After the two distinct cycles, the waves were damped; probably because of their irregularity and variability in the individual cells. Nevertheless, our data indicate that the ribosomal genes are expressed discontinuously, with intervals of 45-75 min between the bursts.

In our review on the discontinuous transcription, we indicated what seemed to be four main patterns in which this phenomenon may be manifested: the typical busts; the undulating pattern; the regular pulsing; and the rare transcription events.[1] As mentioned above, the fluctuations observed in our study do not seem to belong to the regular type. Rare events also must be excluded, since rDNA transcription is very intensive throughout the entire interphase. The typical bursts are separated by the relatively long periods of silence. But we observed no diminishing of the number of FU positive (Fig 7B) or pol I positive (Fig 3) FC/DFC units in the course of the experiment, although the mean intensity of the incorporated FU signal in the individual units was greatly reduced at the points of minimal transcription activity (Fig 7B). Therefore, the observed fluctuation of rDNA transcription most likely belongs to the undulating pattern, in which the bursts are alternated by periods of relatively rare transcription events.

Additionally, our method of synchronization allowed us to obtain averaged data concerning the fluctuations in the nucleoplasmic genes, since their expression was also inhibited by the cold treatment. After this procedure, the total transcription signal in the nucleoplasm showed symptoms of fluctuations with two discernible, though not very distinct, peaks (Fig 6). Evaluating these results, we have to keep in mind that various nucleoplasmic genes in the same cell display a wide range of transcriptional kinetic behavior (reviewed in Smirnov et al. [1]).[4, 10, 55, 56] Moreover, some of these genes are expressed in typical bursts with long periods of silence, during which they cannot be detected by FU incorporation. We should also mention that the status of the nucleoplasmic RNA polymerases at the low temperature was not examined in our experiments, and thus we do not know how efficiently the transcription was synchronized. Nevertheless, the presence of two significant peaks on the periodograms (Fig 6) suggests that numerous genes in the nucleoplasm were transcribed in a pulse-like manner with periods close to 15 min and 75 min.

Thus, our results indicate that ribosomal genes in human cells are expressed discontinuously, and their transcription follows undulating pattern with predominant period of about 60 min.

## Acknowledgments

The work was supported by research project BBMRI_CZ LM2015089, by the Grant Agency of Czech Republic (19-21715S), by research project BBMRI_CZ LM2015089, and by Charles University (Progres Q25 and Q28).

